# Asamataxis: A cooperative relayed migration in response to subsurface inhomogeneity leads to long-range self-patterning of cells

**DOI:** 10.1101/799437

**Authors:** Akshada Khadpekar, Kanksha Mistry, Nehal Dwivedi, Aditya Paspunurwar, Parag Tandaiya, Abhijit Majumder

## Abstract

Cells self-organize to give patterns that are essential for tissue functioning. While the effects of biochemical and mechanical cues are relatively well studied, the role of stiffness inhomogeneity on cellular patterning is unexplored. Using a rigid structure embedded in soft polyacrylamide (PAA) gel, we show that such mechanical inhomogeneity leads to long-range self-organized cellular patterns. Our results reveal that this patterning depends on cellular traction and cell morphology. Depending on a suitable combination of cellular morphology and traction, the information about the presence of embedded structure gets relayed outward. In response to this relay, the cells reorient their axis and migrate towards the embedded structure leading to the observed long-range (20-35 cell length) patterning. To predict the possibility of pattern formation, we present a dimensionless number ‘*f*’ combining the governing parameters. We have also shown that the pattern can be tailor-made by pre-designing sub-surface structures, a potential tool for tissue engineering. This mechanism of directed migration driven long-range pattern formation in response to mechanical inhomogeneity may be involved during several pathophysiological conditions, a proposition that needs further investigation.

**One Sentence Summary:** Substrate inhomogeneity and cooperative cellular traction together lead to cellular migration and long-range pattern formation.

---

Spatial patterning of cells is a common feature in nature and essential for efficient tissue functioning. Understanding the mechanism of multicellular patterning is a key question in developmental biology and tissue engineering. Since 1952, when Alan Turing^1^ proposed reaction-diffusion model in his seminal paper, many diffusion-based patterning mechanisms have been proposed and demonstrated^2^. In recent years, it has been shown that mechanical cues too play important roles in tissue patterning^3–6^. For example, cells align in response to topography^7,8^, shear stress^9,10^, and cyclic strain^11,12^. Cellular alignment can also be controlled by systematically varying the substrate stiffness. For example, Choi et al. showed that when skeletal myoblast C2C12 cells were cultured on substrates with stripes of alternating stiffness, cells were aligned in the direction of the long axis of the stiff regions^13^.

In all these studies investigating the effect of mechanics on cellular patterning, one neglected parameter is mechanical inhomogeneity due to embedded rigid structures inside otherwise soft surroundings. Such situations are not uncommon in physiological/pathological conditions. A few such examples are solid tumors embedded in soft tissue, scars, myocardial infraction, the skeletal structures where rigid bones are surrounded by comparatively soft muscles etc. While the natural examples are plenty, surprisingly, studies to explore the effect of embedded structures on cellular behavior are handful, and on patterning is none. Literature shows that when the rigid object is in close proximity to the surface, the effective rigidity changes. As a result, nearby cells cluster and align on top of the rigid structure, but no long-range patterning was observed. However, what happens when the structure is deeply buried? Can multiple cells together sense the presence of a buried structure that a single cell cannot? Can this information be passed over multiple cells? Moreover, can this rigid body work as an “anchoring point” for the cells to “hold” and align?

To answer these questions, here we present a so far unreported patterning mechanism governed by substrate mechanical inhomogeneity and cooperative interactions of the cells. In this work, we embedded rigid glass objects inside a soft polyacrylamide (PAA) substrate and cultured different adherent cells on this composite substrate. We observed that adherent contractile cells self-organize into a 1-2 mm i.e. ~20 to 35 cell length long pattern in response to this mechanical inhomogeneity. The cells reorient their major axes along the line of maximum resistance from the substrate and migrate towards the embedded rigid object. We name this migration as Asamataxis (“Asama” is unequal or inhomogeneous in Sanskrit). With time, the information is relayed outwards via the deformable substrate and more number of cells take part in this cooperative behavior establishing a long-range pattern. If the cell number is low, then this cooperative mechanism fails, and cells do not show persistent migration towards the bead. Cellular morphology and traction, along with the substrate mechanical property together contribute to this directed migration and pattern formation. We also present an empirical equation to predict the possibility of pattern formation. The pattern can be tailor-made by using pre-designed sub-surface structures, a potential tool to be further explored in tissue engineering.

## Results

### Sub-surface rigid structures induce cellular alignment on the surface of a soft polyacrylamide gel

To understand the effect of mechanical inhomogeneity on cellular behavior, we embedded rigid glass structures into an otherwise deformable polyacrylamide (PAA) gel (Fig. 1A) (details are in method section). For cells to adhere, the gels were coated with extracellular matrix (ECM) protein of choice. For most of our work, we used glass bead of diameter, D = 1.04 ± 0.05 mm (Fig. S1) as our rigid object embedded into PAA gel of stiffness (E)=2 kPa and height (H) = 1.2 mm (Fig. 1A and B), coated with rat tail collagen I. We seeded C2C12 cells, a mouse myoblast cell line, and cultured for 48 h, to understand the effect of mechanical inhomogeneity on cellular behavior. We found that the C2C12 cells self-organize into a radially aligned pattern around the bead (Fig. 1C). In the image, the embedded bead appears black as the images are captured using the phase-contrast microscope. The radial alignment was not observed for cells on homogeneous polyacrylamide gel (E = 2 kPa) without the bead (Fig. 1D) or on regular tissue culture plastic (E ~ GPa) (Fig. 1E). Using OrientationJ plugin of Image J, we further highlighted the global pattern on gel with bead (Fig. 1F). The colors on the map indicate the direction of the predominant cellular orientation with respect to X-axis of image. Fig 1F shows that the orientation of the cells near the bead is correlated with the far away (approx. 1000 µm) cells along the radial line from the center of the bead. However, no such long-range correlation was observed on gel without bead (Fig. 1G) or on plastic (Fig. 1H). Only patches of local cell alignment were present on substrates with uniform rigidities.

**Fig. 1.**
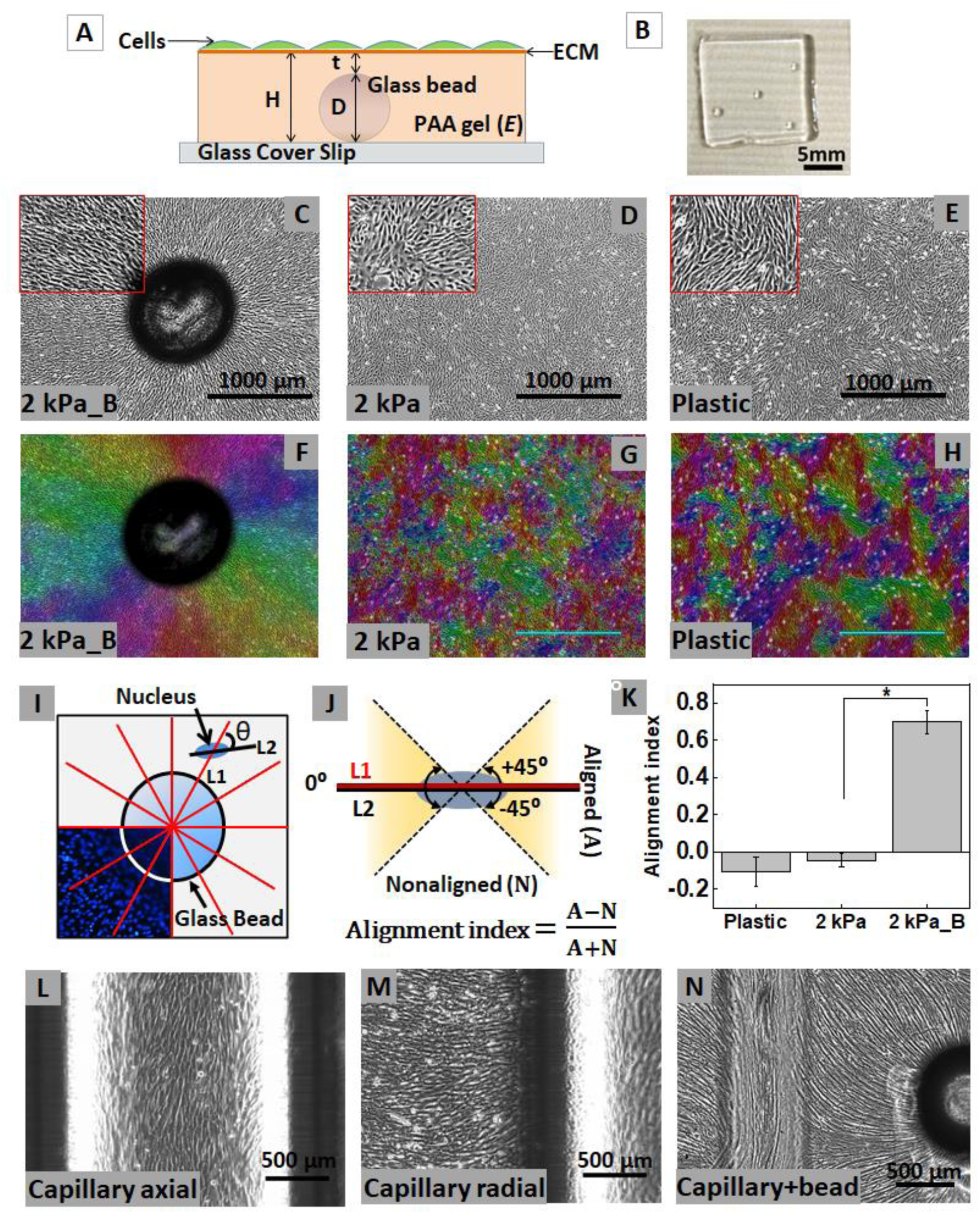
Self-organization of cells due to embedded rigid material in soft PAA gel. (**A**) Schematic of rigid material (here, glass bead) of diameter ‘D’ embedded in polyacrylamide (PAA) gel substrate of height ‘H’ and stiffness ‘E’. The layer of PAA gel above the embedded bead is of thickness ‘t’. This substrate is coated with extracellular matrix (ECM) and seeded with cells. (**B**) Top view photograph of the substrate. (**C**) C2C12, mouse myoblast cells, cultured on this mechanically inhomogeneous substrate self-organize radially around the embedded bead on the soft (2 kPa) PAA gel surface whereas cells on (**D**) 2 kPa PAA gel without bead and (**E**) tissue culture plastic dish and were randomly arranged. (**F-H**) Color-coded map of predominant cellular angle orientations in phase-contrast images of (**F**) the 2 kPa_B, (**G**) TCP and (**H**) 2 kPa gel without bead. (**I**) The radial alignment was quantified as shown in the schematic. The acute angle θ was measured between line L1, drawn from the center of the bead cutting the nucleus and line L2, drawn along the major axis of the nucleus. For analysis of each image, at least 12 such L1 lines were drawn. (**J**) The angle θ with the range between 0⁰±45⁰ was defined as aligned and 90⁰±45⁰ was non-aligned. Alignment index was calculated using the formula shown in (**J**). (**K**) Maximum cells were radially aligned on 2 kPa with embedded bead (2 kPa_B) substrate compared to control (Plastic and 2 kPa only gel). Other than glass bead, C2C12 cells on top of embedded glass capillary aligned axially (**L**) and on sides cells were radial (**M**) to the length of a glass capillary. (**N**) Combination of glass capillary and bead embedded 2 kPa PAA gel also showed alignment of cells. Data is represented as mean ± sem. The total number of cells analyzed from three experiments, n>300.

To quantify the alignment, we stained the nuclei with Hoechst and considered nuclear orientation as representative of cellular orientation (Fig. S2). The angle theta θ between the radial vector starting from the epicenter of the bead (L1) and the major axis of the intersecting nucleus (L2) was used as the measure of radial alignment (Fig. 1I). If θ is between 0⁰ ± 45⁰ (shaded region, Fig. 1J), we call the cell as aligned (A), otherwise non-aligned (N) (Fig. 1J). Further, we calculated the alignment index (AI) using the formula,

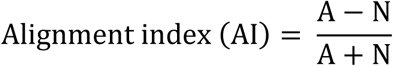

AI approaching +1 and −1 respectively indicates radial and circumferential alignment. AI ~0 signifies unbiased or random alignment for the cell population. We found that for plastic and homogeneous 2 kPa PAA gel, the alignment index was −0.1 ± 0.08 and −0.05 ± 0.03, respectively (Fig. 1K). However, for 2 kPa substrate with bead (**named as 2kPa_B**), AI increased to 0.7 ± 0.10, indicating a strong preference towards radial alignment (Fig. 1K). In other words, on 2 kPa_B substrates, 85% of the cells were radially aligned (Fig. 1K).

### Cellular alignment is dependent on the geometry of the embedded rigid structure

To study the effect of geometry of the embedded structure, we used glass capillary (D=1 mm), combination of beads and capillary, and disks. We found that on top of the capillary, cells were axially aligned (Fig. 1L) whereas, on the side, they were aligned in transverse direction (Fig. 1M). As both the bead and glass capillary had finite curvatures that lead to a gradual change in the substrate thickness (t) on top of the capillary and bead, we asked if this gradual change in (t) is necessary for the cellular alignment. We embedded a disc of stiff (40 kPa) PAA gel inside soft 2 kPa PAA gel substrate keeping the height (H) and thickness (t) unaltered. The cells were aligned radially from the edge of the disc similar to the 2kPa_B substrate (Fig. S3). These observations together suggest that the cellular pattern formation depends on the geometry but a gradual change of skin thickness “t” is not necessary. Furthermore, the patterning can be tailor-made using pre-designed subsurface structures as shown in Fig. 1N, where a combination of glass capillary and glass bead were embedded inside soft PAA gel resulting into a combination of cell patterning.

### Mechanical and geometric parameters of the substrate are important for cellular patterning

To verify, if the mechanism of observed pattern formation is of physical origin, we changed the elastic modulus (E) of the substrate from 2 kPa to 5 kPa, 10 kPa and 20 kPa keeping other parameters unchanged We observed that the radial alignment was gradually lost with increasing (E) (Fig. 2A-E). While on 2 kPa, the cells were radially aligned around the bead (Fig. 1C and 2A, 2A inset), on 5 kPa substrate the cellular radial alignment was somewhat disrupted (Fig. 2B and 2B inset). With further increase in stiffness to 10 kPa and 20 kPa, the long-range radial alignment was completely lost (Fig. 2C-D, and 2C-D inset). The alignment index (AI), as described in Fig. 1J, confirms this visual observation. The AI on 5 kPa substrate was 0.36 ± 0.1 showing a ~50% drop from 0.70 ± 0.10, AI on 2 kPa (Fig. 2E). The AI on 10 kPa and 20 kPa reduces further to ~0 (−0.04 ± 0.12 and −0.01 ± 0.01 respectively), indicating random orientation (Fig. 2E). This observation implies that the increasing substrate stiffness masks the embedded rigid bead making it imperceptible to the cells. Further, we demonstrated that the radial alignment observed was unaffected by changing the ECM from collagen to gelatin and laminin (Fig. S4), highlighting the dominant physical origin of the observed pattern.

**Fig. 2.**
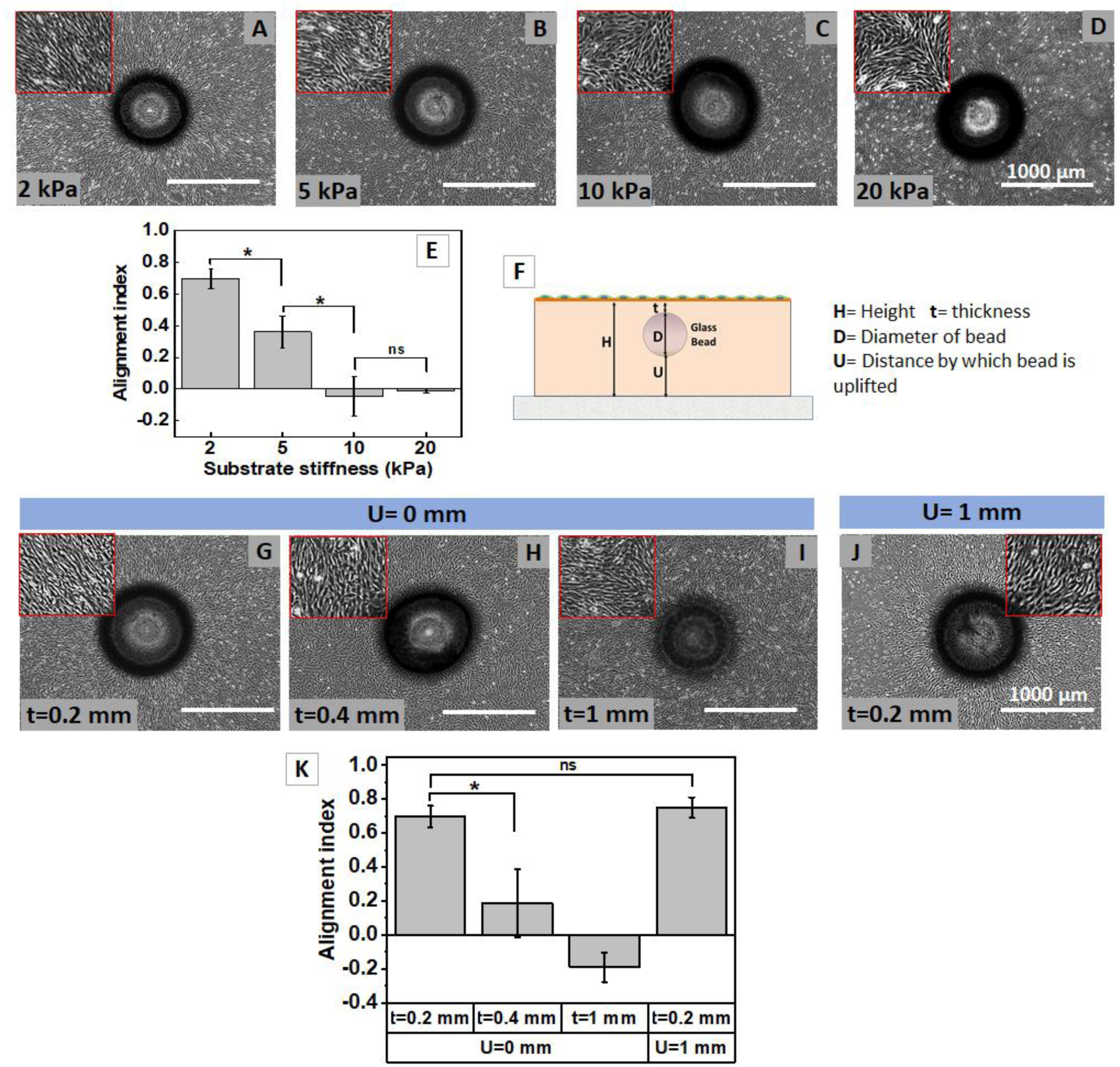
Cell patterning is influenced by the mechanical and geometrical parameters of the substrate. (**A-D**) Phase-contrast image of C2C12, myoblast on glass bead embedded PAA gel of stiffness (**A**) 2 kPa, (**B**) 5 kPa, (**C**) 10 kPa and (**D**) 20 kPa. (**E**) The alignment index approached zero with increasing stiffness which indicates that the radial alignment is lost with increasing stiffness. (**F-I**) The increase in substrate thickness ‘t’ from (**G**) 0.2 mm where we see radial alignment of cells to (**H**) 0.4 mm and (**I**) 1 mm, there is a loss of radial alignment pattern (**K**). (**F**) The schematic of uplifted bead substrate to decouple between ‘H’ and ‘t’. The bead was uplifted ‘U’ by 1 mm in the PAA gel substrate with H=2.2 mm, D=1 mm so that ‘t’ is 0.2 mm. (**J-K**) The radial alignment was restored on the uplifted bead substrate. Data is represented as mean ± sem. The total number of cells analyzed from three experiments/beads, n>300.

Next, to check the effect of substrate geometry, we increased substrate height (H) from 1.2 mm to 1.4 mm and 2 mm, keeping substrate elasticity (E) and bead diameter (D) constant. However, to note, in this experimental design, gel height (H) and thickness (t) were not mutually independent parameters. For example, when we changed the substrate height from 1.2 mm to 2.2 mm, the thickness (t) correspondingly changed from 0.2 mm to 1.2 mm. The cells were patterned on 0.2 mm (Fig. 2G) PAA gel thickness but failed to organize radially when thickness (t) was increased to 0.4 mm (Fig. 2H) and 1 mm (Fig. 2I). The AI values for substrate with t = 0.4 mm and t= 1 mm were 0.19 ± 0.20 and −0.19 ± 0.08, respectively (Fig. 2K). To decouple (H) and (t), we raised the bead by a distance U = 1 mm from the base glass coverslip in a gel of height H = 2.2 mm (Fig. 2F). This modification helped us to maintain t = 0.2 mm with increased H. The lifting of bead restored the radial alignment, further confirming the role of mechanics in this process (Fig. 2J-K). The AI value for cells on the substrate with raised bead (U=1 mm) was 0.75 ± 0.06, similar to that on 2 kPa_B substrate (t =0.2 mm and U=0 mm) (Fig. 2K). To summarize, the stiffness of the substrates and the depth of the bead from the gel surface, both are important for pattern formation. However, it is important to note that the depth, i.e. 200 µm, for which we report bead sensing here is beyond the reported values of depth-sensing in literature^14,15^

### Embedded structure establishes pattern ranging ~20-35 cell length

As we have observed in previous sections, cells get radially aligned in response to the embedded bead and form a pattern over a long-range (Fig. 3A and B). To find the range of this multi-cellular pattern, the alignment angles (AA) of cells were measured as described in Fig. 1I and J. The area around the bead was divided into multiple annular regions, each of 100 µm width. The average AA (AA_average_) for all cells situated within a particular annular region was estimated and plotted against the distance of that annular region from edge of the bead (Fig. 3C). The farthest distance till which this AA_average_ <45° is defined as the range of cellular alignment. Based on this analysis, we found that for D = 1 mm (corresponding H = 1.2 mm, and t = 0.2 mm the range of alignment was 1100 µm from the perimeter of the bead as marked with yellow dotted line in Fig. 3A. Considering the average length of an elongated cell is ~50 µm, the observed pattern ranges over more than 20 cell length. This range increased to 1800 µm (~35 cell length) when the diameter of the bead was doubled to D = 2 mm, keeping (t) unchanged (Fig. 3B and C).

**Fig. 3.**
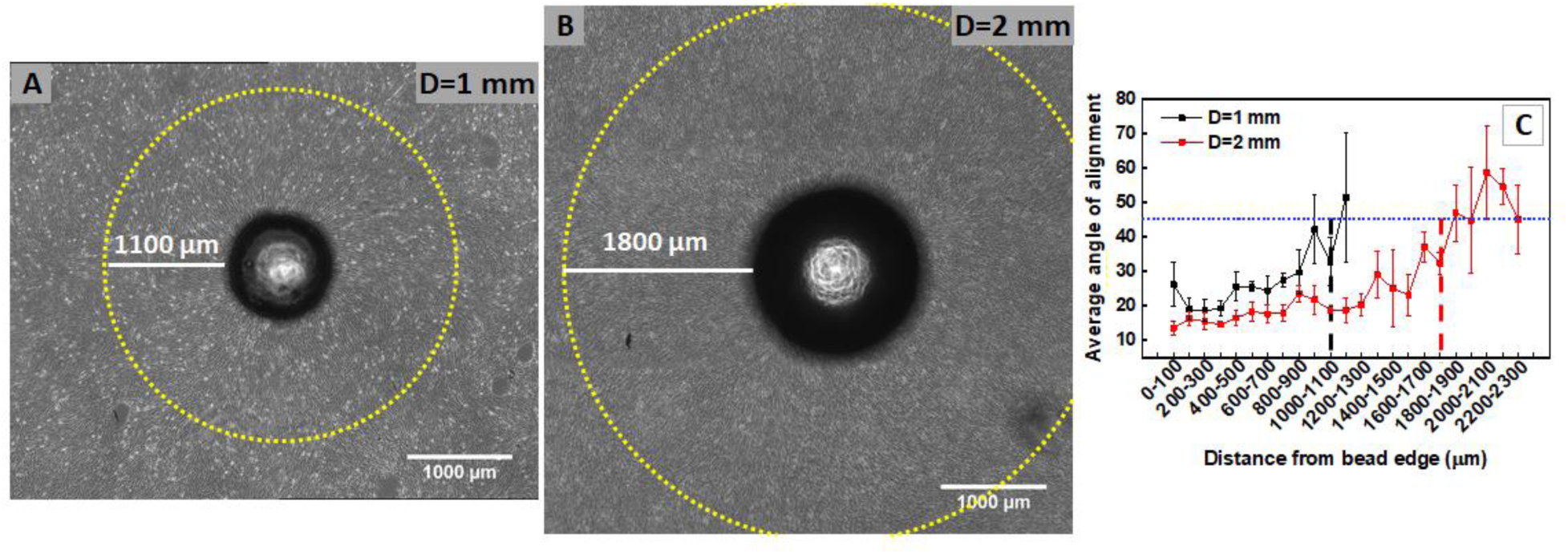
Long-range cell alignment depends on the diameter of the embedded bead. (**A**) On substrate with H=1.2 mm, t=0.2 mm and D=1 mm, the range of radial cell alignment from the perimeter of the bead is 1100 µm (yellow circle). (**B**) When the bead diameter was increased to 2 mm, with constant t=0.2 mm, the cell alignment increased to 1800 µm (yellow circle). (**C**) The average angle of alignment plotted against distance from the bead periphery. The cut off of 45° was used to differentiate between aligned and non-aligned cells. The distance just before the one at which the average angle is more than 45° is the range of radial alignment. Using this, for substrate with D=1 mm, the range of aligned cells is 1100 µm as marked by black dotted line while for substrate with D=2 mm, it was 1800 µm (**C**). The data is plotted as mean ± sem. (n>400 cells for D=1 mm and n>750 cells for D=2 mm, N=3)

### Biased cell migration towards embedded rigid structure in soft PAA gel

Next, we studied whether this long-range cell alignment is due to the transmission of mechanical force. We specifically asked three questions; 1. Whether the participating objects, here cells, show a cooperative directed migration, 2. What is the Spatio-temporal dynamics of this self-organization, and 3. How the information is passed over the observed range which is two orders of magnitude higher than the cellular length scale. To answer these questions, we captured time-lapse images with 30 min interval for 48 h on soft (2 kPa_B) (Video S1) and stiff (20 kPa_B) (Video S2) PAA gel.

For each cell, we drew the displacement vector R as |R1|-|R2|, where R1 and R2 are the initial and the final position vectors originating from the epicenter of the bead, as shown in schematic Fig. 4A. Representative images of such displacement vectors (R) for all the cells tracked within the field are shown in Fig. 4B (for 2 kPa) and Fig. 4C (for 20 kPa). The colored arrows in the image are the displacement of each cell. The bead periphery is shown as a white dotted circle. Arrow tail (the filled circle) and arrow head indicate the initial and final position of a cell, respectively. On 2 kPa_B substrate, maximum arrows were pointing towards the bead/white dotted circle whereas on 20 kPa_B, the arrows were pointing in random directions. Further, if |R1|-|R2|>0, we defined the cell to be migrating towards the bead, else migrating away. We found that on soft gel, as high as 75% of the cells were migrating towards the bead as three quartiles of the box plot of |R1|-|R2| was greater than zero (Fig. 4D). No such bias was observed on 20 kPa substrate, where |R1|-|R2| values were equally distributed around zero (Fig. 4D). We name this so far unreported cooperative cellular migration as Asamataxis. We eliminated the possibilities of haptotaxis and durotaxis by confirming uniform collagen density and constant rigidity of the gel using fluorescence microscopy and AFM, respectively (Fig. S5).

**Fig. 4.**
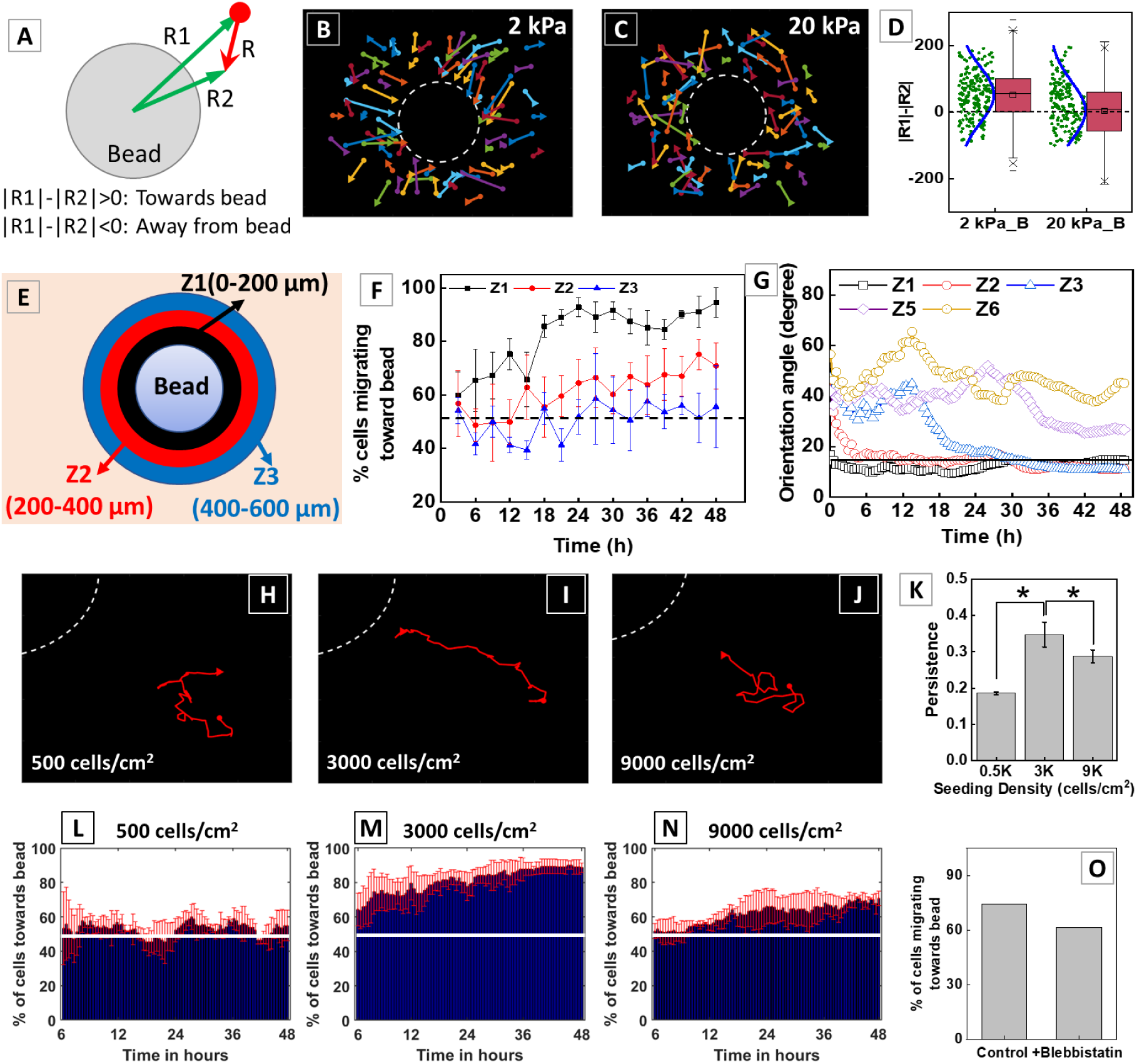
Relay of mechanical signal from bead periphery to far away cells is required for cooperative migration and long-range pattern formation. (**A**) The schematic of the method used to calculate cell migration towards and away from the bead. Representative plots of cell displacement after 48 h of cell seeding on (**B**) 2 kPa (n= 75 cells) and (**C**) 20 kPa (n=56 cells) with embedded bead substrates. The colored arrows indicate the cell displacement and the white dotted circle is the perimeter of the embedded bead. (**D**) On 2 kPa with embedded bead substrate, maximum cells have positive R1-R2 which indicates biased cell migration towards bead (n>200). (**E**) The schematic showing the image divided into zones from the bead center, Z1 (0-200 µm), Z2 (200-400 µm) and Z3 (400-600 µm) for to determine the relay. (**F**) The % of C2C12 migrating towards bead on 2 kPa_B from each zone was calculated and found that cell in Z1 migrates first towards bead followed by cells in Z2 and the cells in Z3. The data is plotted as mean ± sem (n>150 cells, N=3). (**G**) The orientation angle of cells <15° (black line) are aligned. Cells in Z1 are aligned from 0.5 h while cells in Z2 and Z3 were aligned after 11.5 h and 30 h, respectively. The cells in Z5 and Z6 were never aligned radially around the embedded bead. (Note: Z4 is not shown to avoid crowding. The cells in Z4 were aligned after 38 h.) (**H-J**) The representative cell track in a (**H**) low (500 cells/cm^2^), (**I**) medium seeding (3000 cells/cm^2^) and (**J**) high seeding (9000 cells/cm^2^) density migrating towards bead. (**K**) The persistence of cells at low and high seeding density was less compared to cells at medium seeding density. The white dotted curve (**H-J**) is the periphery of the bead. (**L-N**) Similar to persistence, the number of cells migrating towards embedded bead at (**L**) low, (**M**) medium and (**N**) high seeding density was 50%, 90% and 75%, respectively. (**O**) The percentage of cells migrating towards bead decreases with 20 µM blebbistatin treatment (n=40 cells).

### Mechanical signal relays out from embedded rigid structure to far away cells

Earlier, we have shown that the range of pattern formation is upto 1100 µm from the bead periphery (Fig. 3). It is unintuitive how a cell 1000 µm away from the bead can sense the inhomogeneity. To solve this puzzle, we first explored the spatio-temporal dynamics of the biased cell migration. For this, we divided the area around the bead into multiple annular concentric zones each of 200 µm width, so that, zone 1 (Z1, black) = 0-200, zone 2 (Z2, red) = 200-400, zone 3 (Z3, blue) = 400-600 µm, and so on (Fig. 4E). We calculated the percentage of cells migrating towards the bead for each zone in every 3 h intervals. We found that from very early time point, 60% of the cells in Z1 were migrating towards the bead. Migration bias increased steadily for the first 24 h to finally reach a plateau value of 90% which was maintained for the rest of the duration of our observation (Fig. 4F). Cells in Z2 lagged behind Z1 and was unbiased for first 12-18 h. After that, the percentage of cells migrating towards the bead in Z2 increased steadily to reach 75% at the end of our observation time (48 h). The percentage of cells migrating towards bead in Z3 was always close to 50%, indicating unbiased migration (Fig. 4F). Estimation of the cellular alignment in a similar spatio-temporal manner confirms that the formation of radial pattern also follows the same relay mechanism. We observed that the cells in Z1 were aligned after 0.5 h of cell seeding, while cells in Z2 and Z3 were aligned after 11.5 h and 30 h, respectively (Fig. 4G). In Z5, although the quality of alignment was not similar to that in previous zone, orientation angle decreased after 36 h and then plateaued. Interestingly, cells in Z6 (1000-1200 µm) were not aligned at all during our observation time. This confirms that along with the biased cell migration, cell alignment also relays from inner zones to outer zones with time which indicates that the cells near the bead sense the inhomogeneity first and the information relays outwards with time.

### Neighboring cells influence persistence of cells migrating towards embedded bead

To check how cell crowding influence the directed migration, we seeded C2C12 cells at varying seeding density of 500 cells/cm^2^ (low) (Video S3), 3000 cells/cm^2^, (medium) (Video S4) and 9000 cells/cm^2^ (high) (Video S1) on 2 kPa_B substrate. The representative cell tracks at low (Fig. 4H), medium (Fig. 4I) and high (Fig. 4J) are shown in red color in Fig. 4H-J respectively. The white color dotted line indicates the embedded bead periphery. At medium seeding density, cells showed significantly higher persistence compared to the same at low and high seeding density (Fig. 4K). This suggests an optimum cell number is needed to establish the biased migration in response to the substrate inhomogeneity. If the cell number is low, there is not enough cells to relay the message outwards. As a result, percentage of cells migrating towards the bead (Fig. 4L-N) as well as the migration persistence decreases (Fig. 4K). On the other hand, for high seeding density, by the time, the information reaches farther zones from bead periphery, the zone becomes crowded due to cell division restricting the scope of directed migration, as was observed in Fig. 4K. Also, this indicates that for directed migration to happen we need some optimum initial cell density near the bead.

Cell migration and alignment are known to depend on cellular contractility^16,11^. In this work too, when we perturbed the cellular contractility by inhibiting the myosin using blebbistatin, the biased migration of cells towards embedded bead on soft substrate was lost upon treatment with blebbistatin (Fig 4O). Together, we show that the cells at an optimum distance from each other applying traction force on the soft deformable substrate form the long-range pattern via mechanical relay.

### Sustained cellular traction stress is essential for cell patterning

In Fig. 4O, we have shown that the biased migration is lost when the cells are treated with traction inhibitors. Also, the radial alignment was lost on inhibiting the traction stress using blebbistatin and nocodazole (Fig. S6). Here, we ask the reverse question, can we form the pattern by increasing the cellular traction in a situation where usually the pattern is not formed? To address this question, we treated the cells on 5 kPa_B substrates, with LPA (30 µM), a molecule known to enhance the cellular traction via Rho activation^17^. We found that with an increase in cellular traction, AI increases from 0.36 ± 0.09 (without LPA/control) to 0.64 ± 0.10 (with LPA) (Fig. 5A-C).

**Fig. 5.**
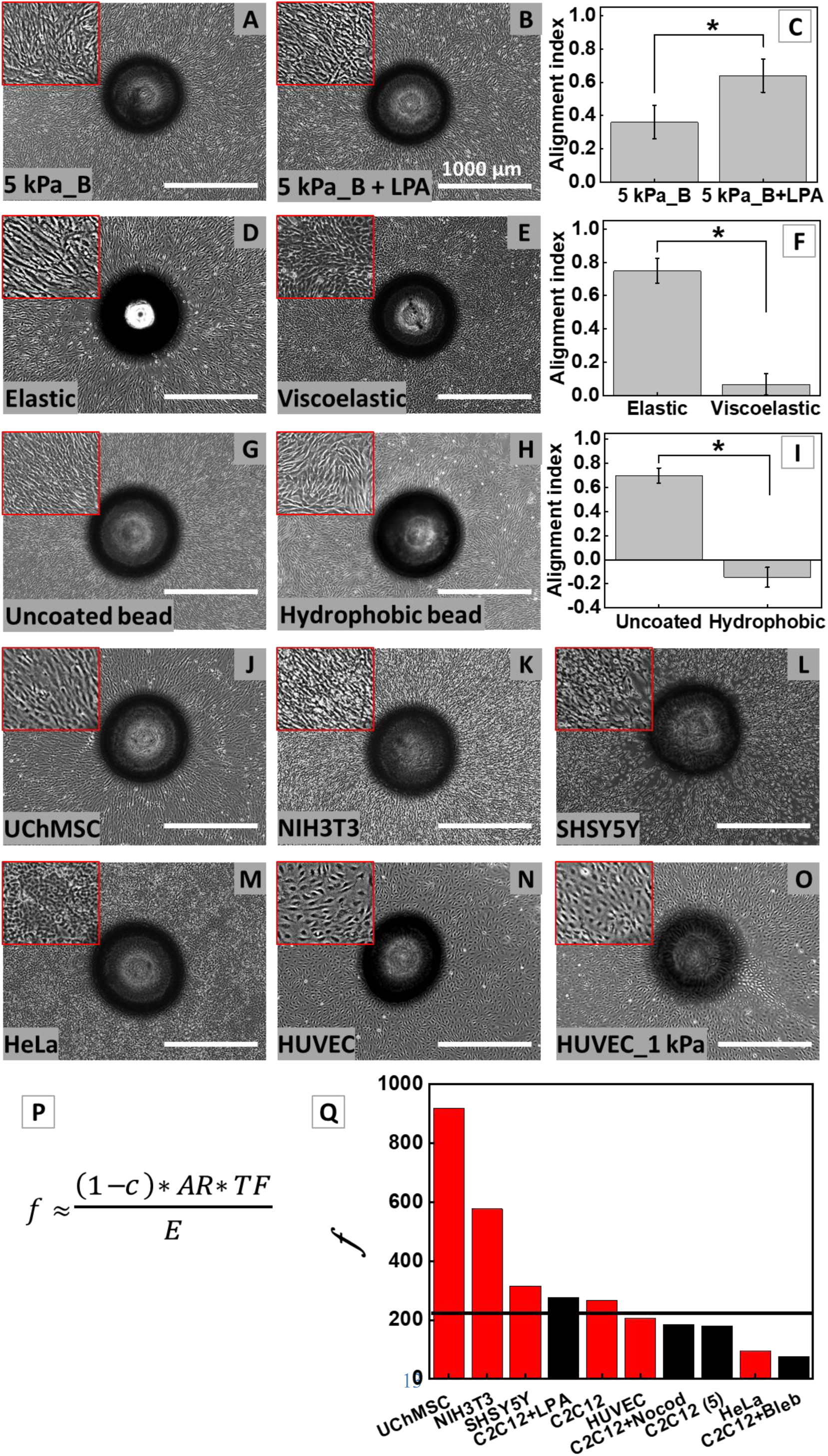
Traction force and cellular morphology-players in cellular patterning. (**A-C**) Increase in traction force leads to pattern formation. The C2C12 cells on (**A**) 5 kPa PAA gel with bead (5 kPa_B) were partially aligned. However, the cells treated with LPA (30 µM) on 5 kPa_B substrate showed radial alignment (**B**). (**C**) The alignment index increased significantly on 5kPa_B substrate treated with LPA. (**D-F**) Sustained traction is required for pattern formation. Phase-contrast image of C2C12 cells on (**D**) elastic and (**E**) viscoelastic PAA gel substrate with embedded bead. (**F**) The cells on viscoelastic substrate were randomly oriented compared to elastic substrate with embedded bead. (**G-I**) Bead gel interaction is required for pattern formation. Phase-contrast image of C2C12 on the (**G**) uncoated bead embedded substrate and (**H**) hydrophobic (DCDMS) solution coated glass bead embedded in 2 kPa PAA gel. (**I**) The AI value of cells drastically dropped when the bead-gel adhesion is disrupted. (*p<0.05, n>250 cells, N=3). Phase contrast image of (**J**) UChMSC, (**K**) NIH3T3, (**L**) SHSY5Y cells aligned on 2 kPa_B substrate whereas (**M**) Hela and (**N**) HUVEC cells were not aligned radially. (**O**) Phase contrast image of HUVEC cells aligned on 1 kPa_B substrate. (**P**) A dimensionless parameter ‘*f*’ which is used to predict the cellular pattern formation. (**Q**) The values of ‘*f*’ are plotted for different cell types and conditions. The conditions or cell where cells were not aligned have value <207. The column in red color indicates different cell types and the black color bar indicates inhibitor conditions. (n>15, N=1)

Further, to verify the essentiality of sustained traction, we first used viscoelastic PAA gel where traction stress is known to decay with time^18^ and compared with the pattern formed on elastic gel with similar storage modulus. We used elastic and viscoelastic with same storage modulus (G’) = 1.2 kPa and different loss modulus (G”) = 30 Pa and 300 Pa, respectively. We found that while the cells aligned radially on elastic gel, they fail to form the pattern on viscoelastic gel of same rigidity (Fig. 5D-F). The AI was found to be 0.75 ± 0.07 and 0.07 ± 0.06, for elastic and viscoelastic gels, respectively (Fig. 5F).

Next, we ask if the bonding between the embedded structure and the gel is essential for pattern formation. We found that if we allow the gel to relax/slip at the bead surface by chemically treating the glass bead to reduce the bead-gel adhesion, the cellular pattern fails to form (AI = −0.15 ± 0.08, Fig. 5G-I). These results together confirm that that sustenance of traction stress and resistance from the bead via gel-bead bonding is essential for the cellular pattern formation.

### Pattern formation is cell type-dependent

Next, we asked the question what if we use different cell types which differ substantially in their shape, size and traction. To answer, along with C2C12 that we have already described throughout this paper, here we used Umbilical cord human mesenchymal stem cells (UChMSCs), terminally differentiated mouse fibroblast (NIH3T3), human neuroblastoma (SHSY5Y), human cervical cancer epithelial cells (HeLa) and human umbilical vein endothelial cells (HUVEC) on 2 kPa_B substrate. Out of these cells, C2C12, UChMSCs, NIH3T3 and SHSY5Y cells showed radial alignment (Fig. 5J-L) but HeLa and HUVEC cells did not (Fig 5M-N). However, HUVEC cells could radially align on 1 kPa_B substrate (Fig. 5O), HeLa did not pattern even on 1 kPa_B (Fig. S7).

### Traction force and cellular morphology together control the pattern formation

To understand the origin of cell-dependent variability, we estimated the traction of each of the cell type mentioned above. Interestingly, we found that while C2C12, NIH3T3 and HUVEC cells have similar traction, C2C12 and NIH3T3 can form patterns but HUVEC cannot. In contrary, SHSY5Y cells have very low traction yet capable of forming long-range patterns. We hypothesize that the neuronal cells can overcome the limitation of low traction by measuring the stiffness at distant points due to their high aspect ratio. This shows that the cellular traction alone cannot explain the success/failure in pattern formation in these cells (See the traction and relevant morphometric parameters in Table. S1). We further noticed that SHSY5Y cells have low traction but high aspect ratio. As spindle-shaped cells concentrate their traction stress at their two ends^19^, we hypothesize that this characteristic enables the cells to efficiently relay the mechanical information to their neighbors compared to their low aspect ratio counterparts. Also, the cell with sharp edges applies more concentrated traction compared to smooth edge cell of the same size and aspect ratio^20^. As a result, the cells with high aspect ratio (AR) and low circularity (c) can form long-range patterns even with low traction. Combining these parameters along with the substrate elasticity (E), we propose a dimensionless parameter ‘*f*’ to predict the alignment as shown in Fig. 5P. Introducing circularity in the numerator in the form of (1-c) forces the parameter ‘*f*’ to approach zero for a circular cell. Such arrangement for ‘c’ is reasonable because a circular cell cannot form a closed pack radial pattern.

The values of ‘*f*’ for different cell types were found to be ~919, 577, 315, 267, 207 and 95 for UChMSC, NIH 3T3, SHSY5Y, C2C12, HUVEC and Hela, respectively (Fig. 5Q). To note, all the cell types that showed alignment had corresponding ‘*f*’ to be greater than 210. The values of the ‘*f*’ was low for HeLa (f=95) and HUVEC (f=207) cells which were not aligned (Fig. 5P). The lowest value of ‘*f*’ for aligned cells was 267 for C2C12 and the highest value for non-aligned cell was 207 for HUVEC cells. To verify the importance of ‘*f*’ in predicting alignment, we perturbed the parameters by treating C2C12 with 20 µM blebbistatin, 0.25 µM nocodazole for 24 h on 2 kPa PAA gel. On treatment with both the inhibitors, the radial alignment was lost (Fig. S6). With the blebbistatin treatment, TF and circularity were reduced by almost 10- and 5-folds, respectively. However, the AR remained unchanged. Whereas, with nocodazole treatment, there was a significant increase in both the TF and circularity, while AR was decreased. The value of ‘*f*’ for blebbistatin and nocodazole treated C2C12 cells on 2 kPa PAA gel was 76 and 185, respectively. Both the values are <210, which supports the aforementioned observation that *f* >210 is for cellular alignment. We have shown previously that the cells on 5 kPa_B substrate are partially aligned (Fig. 2B). The value of ‘*f*’ for C2C12 on 5 kPa PAA gel is 178.7, which further supports our parameter ‘*f*’. To summarize, all the ‘*f*’ values for nine conditions, which includes six different cell type, two inhibitors treated C2C12 and one change in elastic modulus of substrate, shows that there exists a threshold (*f* =207 to 267) for the dimensionless number *f*, above which we get the alignment and below which we do not see any alignment (Fig. 5Q).

### Finite Element Analysis

We have shown that the information about presence of the bead gets relayed outwards resulting in the observed pattern formation. However, the question remains how the first line of cells closest to the bead sense the embedded bead before the information gets relayed. Compared to the earlier literature, the thickness of the gel on top of the bead in our system is 10 times thicker than the reported critical thickness. Hence, it is not surprising that AFM measurement did not show presence of any stiffness gradient (Fig. S5). To investigate if there is a small gradient which comes within the noise of AFM measurement, three-dimensional Finite Element Analyses were conducted. A typical Finite Element Mesh of half the model cut by an X-section passing through the center of the glass bead is shown in (Fig. 9). The details of the FEM model can be found in Materials and Methods section. Here, we are interested in calculating the apparent stiffness of the top surface of the PAA gel substrate as a function of the distance across the embedded glass bead. A displacement of 20 μm is applied on a node on this path in either Y or Z direction, and the resultant apparent stiffness of that point in Y or Z direction was computed. This is referred to as apparent tangential or normal stiffness.

Fig. 6A shows the variation of normalized apparent normal stiffness with distance across the embedded glass bead on the top surface of the PAA gel. In these plots, the glass bead is located at zero position. The apparent stiffness has been normalized by the stiffness value at 2000 μm distance away from the glass bead center, where effect of the bead is not felt. Fig. 6A shows that normalized apparent stiffness exhibit a smoothly varying gradient towards the bead center. The gradient is steeper for small t = 100 μm and progressively reduces to almost zero for large t = 1000 μm. Thus, it can be concluded that for smaller t values, the embedded glass bead exerts more influence on the apparent stiffness of the PAA gel substrate surface. This analysis supports our observation that the effect of bead on cellular pattern formation disappears with increasing ‘t’ (Fig. 2K).

**Fig. 6.**
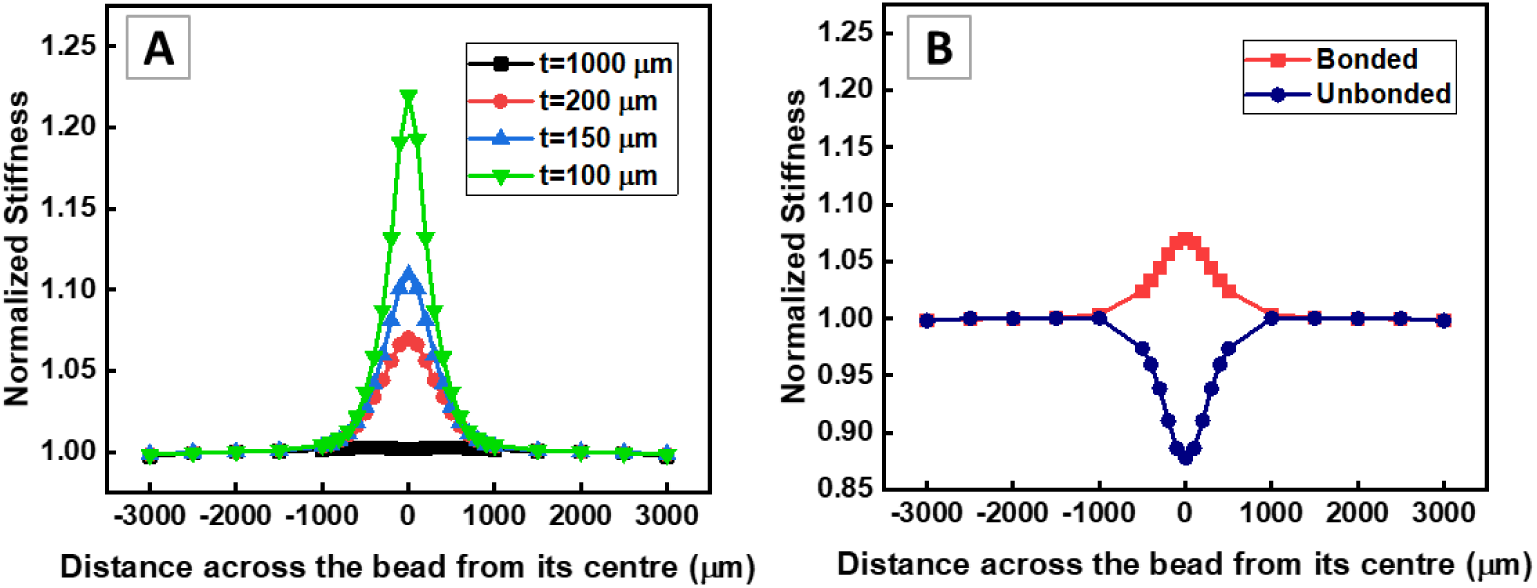
FEM analysis to determine stiffness variation across bead. Effect of (**A**) thickness ‘t’ of the PAA gel above the embedded glass bead and (**B**) interface condition on the variation of normalized apparent stiffness versus distance across the bead.

Further, we examined the effect of interface condition between the glass bead and the PAA gel on the normalized apparent stiffness versus distance across the bead (Fig. 6B). It can be noticed that when the interface is perfectly bonded, an apparent stiffness gradient exists near the bead. On the other hand, when the interface is unbonded and frictionless, the magnitude of apparent stiffness gradient reduces and changes its sign. A minima in apparent stiffness is observed at the bead center for the unbonded interface case. This effect is because shear and tensile tractions cannot be transmitted across the unbonded bead interface. This observation aids to explain our earlier observation made in Fig. 5 H-I.

Together, our FEM analysis shows that the bead increases the apparent stiffness of the gel. For t = 200 µm, as mostly used in this paper, this increase is ~10% at the bead center.

## Discussion

Altogether, this paper presents a novel mechanism of long-range multicellular pattern formation in response to mechanical inhomogeneity. The patterning process is driven by the relay of cellular traction force transmitted via soft gel. There are handful of literature that prepared mechanically inhomogeneous substrates by embedding rigid structure in a compliant material and studied the effect on cellular behaviour^13,14,21–23^. In all these studies, the layer of compliant material above the embedded rigid structures was significantly thin (<20 µm). Due to which the apparent stiffness of the compliant material increased, creating a stiffness gradient on the surface. The cells can sense this apparent difference in rigidity and migrate towards the stiffer region, similar to durotaxis^16^. Due to such migration, cells cluster on the rigid region and forms pattern. However, such patterns are only limited to the dimension of the embedded structure^13^, unlike shown in this manuscript.

If the thickness of the top layer was more than the critical limit (15 µm), the cells could not sense the underlying stiff material^14^. However, in our study, the extent of depth-sensing is far beyond this critical limit. We showed that the cells self-organized even when the top layer thickness was 200 µm. As the top layer was thick in our study, any change in apparent stiffness was non-perceptible by AFM measurements (Fig. S5C-D). However, our FEM analysis captured a 10% increase of apparent stiffness at top of the bead (Fig 6A). At the bead periphery, the percentage increase in apparent stiffness reduces to ~3%.

This difference in the critical thickness of depth-sensing between our study and the literature emerges from the fact that all the previous studies were done using sparsely seeded cells while we used much higher cell densities. The increase in cell number increases the range of mechanosensing via collective cellular traction force. In our previous study, we have shown that the range of horizontal deformation of soft PAA gel increases with the increasing number of cells^24^. Another study by Tusan et. al. showed that the cell colonies can sense the underlying rigid material through a 200 µm thick soft gel, the thickness through which a single cell cannot sense the underlying rigid material^25^. However, while they used cell colonies, we showed that single cells in close vicinity to each other can sense the embedded rigid structure from far away distance due to relay of the strain field via collective traction.

With the above explanation, it is now evident that the embedded rigid structure (here glass bead) acts as an anchor such that the lateral deformation of PAA gel due to collective cellular traction is constrained at the PAA gel-embedded bead interface. This lateral constraint renders the cells near the periphery of embedded rigid structure to overestimate the stiffness of the PAA gel and align their strain axis accordingly (radially around bead periphery). However, it was still not intuitive to explain how a cell >500 µm away can sense the embedded rigid structure. We have shown that the mechanical information is relayed from the embedded rigid structure to the far away cells. The cells near the bead are first to sense the bead. They relay this information in the form of local deformation of the soft gel to cells in the neighboring zones causing them to migrate towards the embedded rigid structure (Fig. 4B) resulting into a long-range pattern formation.

This suggested mechanism is supported by our observations that the pattern is lost 1. with increasing substrate stiffness (E) (Fig. 2A-2E) and thickness (t) (Fig. 2F-2G, 2J), 2. when the adhesion between PAA gel and embedded bead was disrupted (Fig. 5G-5I), 3. when the nature of PAA gel was changed from elastic to viscoelastic (Fig. 5D-5F), and 4. when the cellular traction is disrupted (Fig. S6). As the stiffness of the PAA gel increases the zone of deformation of PAA gel due to force applied by cells decreases^26,27^. Also, the depth sensing ability of collective cell is less than 200 µm^25^. Therefore, when the stiffness of PAA gel or top layer of PAA gel was increased, the patterning disappeared. Also, when the gel-bead interaction is disrupted, the deformation of PAA gel is no longer resisted by the embedded bead and hence the cells behave as if on the homogeneous soft PAA gel giving rise to random cell orientation (Fig. 1D and 1G). The loss of pattern on the PAA gel of viscoelastic nature because the stress field formed by the cellular traction force dissipates with time due to the stress relaxation behavior of the viscous gel. When the traction force is reduced by inhibiting acto-myosin contractility the pattern is lost whereas if the cellular traction was increased on 5 kPa_B substrate, the pattern was restored (Fig. 5A-5C).

In our previous study, we showed that cells form network when seeded on soft substrate at a density such that the average inter-cellular distance is less than 150 µm^24^. Similar global network patterns were shown by Reinhert-King et al.^27^. They showed that when two adherent cells are close enough on a soft deformable substrate, they reorient themselves to align their strain axis forming network patterns. Similar observations were also made computationally^28^. However, in all these studies there was no control over the pattern emerged due to cell-cell interaction. In our study, we demonstrated that controlled cellular pattern can be generated by predesigning the embedded rigid structure (Fig. 1L-1N). We obtained radial sunflower like pattern originating from the epicenter of an embedded glass bead in soft PAA gel (Fig. 1C). When we changed the geometry of the embedded rigid structure the pattern formation on the surface also changed accordingly. The cells on top of the embedded glass capillary were aligned axially while the cells on both sides of the capillary were aligned in perpendicular to the capillary length. For embedded array of beads, we obtained the cellular patterns similar to the magnetic lines observed between two magnets (Data not shown). By embedding different combinations of such structures and playing with substrate stiffness, many different patterns can be created.

The implications of these work can be many. For example, in the neo-angiogenesis during cancer development, the tumor cells secret VEGF which drives the endothelial cell to migrate towards the tumor. However, the stiffness of solid tumor is higher in comparison to surrounding soft tissue which may lead to migration of endothelial cells towards tumor using the mechanism explained in this work. The endothelial cells far away from solid tumor may be able to sense the tumor by cooperative relay of mechanical signal from tumor to the blood vessel. Such possibilities demand further investigation.

In conclusion, we developed a mechanically inhomogeneous PAA gel substrate to study pattern formation. Using this system, we were able to pattern the cells by predefining the geometry of embedded rigid structure. Further, we showed that this long-range patterns were due to sustained cellular traction force and can be relayed to far away cells.

## Materials and Methods

### Preparation of rigid structure embedded polyacrylamide gel substrates

Polyacrylamide (PAA) gels solutions of known stiffness were prepared according to Justin Tse et al.^29^. Briefly, Acrylamide (Sigma-Aldrich) solution was mixed with bisacrylamide (Sigma-Aldrich) solution at different volume. The volume was make up with milli-Q water. Polymerization was initiated with 1/100th total volume of 10% Ammoniumpersulphate (APS) (Sigma-Aldrich) solution and the 1/1000th total volume of N,N,N,N-Tetramethylethylenediamine (TEMED) (Sigma-Aldrich).

Glass bead embedded PAA gel substrates of controlled thickness were prepared as shown in Fig. 7A. The bubble free mixture of poly(dimethylsiloxane) (PDMS) (DowCorning, Sylgard 184) pre-polymer and cross-linker (10:1) was poured on the di-chloro-dimethyl silane (DCDMS) (Sigma) pre-treated glass slides with spacers of required thickness. Another DCDMS pre-treated glass slide was placed on top of this to make a sandwich of the PDMS solution. This PDMS was cured for 24 h at 60°C and then the elastic PDMS layer was used to make the window of 20 x 20 mm dimension. This PDMS window was placed on 3-(Mercaptopropyl) trimethoxysilane (MPTS) (Sigma-Aldrich) functionalized 22 x 22 mm coverslip. 2-3 thoroughly cleaned glass beads of 1 mm or 2 mm diameter were placed inside this window and the polymerizing PAA gel solution of desired stiffness was poured in this window. The DCDMS treated slide was placed on the polymerizing PAA gel solution and pressed slightly to remove any excess solution. This substrate sandwich was allowed to polymerize for 10-15 min after which the top slide and the PDMS window were removed. The glass bead embedded in PAA gel substrate was thus prepared. The top view photograph of actual substrate is shown in Fig. 1B. After gelation, the substrates were soaked in 70% ethanol for 1 min and washed with DPBS for 3-4 times. For all the substrates, if not otherwise specified, height H = 1.2 mm, bead diameter D = 1 mm, and Young’s modulus E = 2 kPa.

**Fig. 7.**
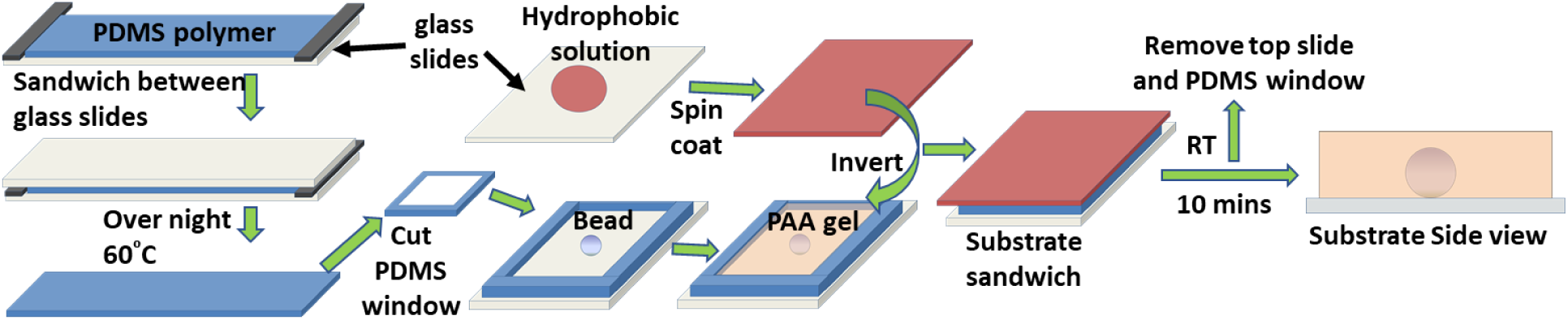
Schematic of rigid structure embedded polyacrylamide (PAA) gel substrate. Schematic for the preparation of mechanical inhomogeneous substrate i.e. rigid structure embedded in polyacrylamide gel substrate.

For gel and bead adhesion study, the beads were coated with dichlorodimethylsilane (DCDMS) for 30 min to make it hydrophobic and then used for substrate preparation.

The viscous gel was prepared using 8% acrylamide and 0.26% bisacrylamide solution and the corresponding control was prepared using 11.4% acrylamide and 0.06% bisacrylamide solution.

### ECM coating on the substrates

The substrates were coated with 100 µl of 5 µg/ml N-sulfosuccinimidyl-6-(4azido-2-nitrophenyl amino) hexanoate (Sulfo-SANPAH) (G-Bioscience) in HEPES buffer (pH=8.5) using UV cross-linker (Genetix) (312 nm) for 20 min. PAA gel substrates were then washed with DPBS twice to remove free molecules of sulfo-SANPAH and submerged in the gelatin (0.1%) or collagen (50 µg/ml) or fibronectin (25 µg/ml) solution and incubated at 4°C for overnight. Finally, before cell seeding, the substrates were kept in laminar hood under UV for 20 min following the washing with DPBS. The substrate is now ready for the cell culture. The schematic of cells seeded substrate is shown in Fig. 1A.

### Cell Seeding on Substrates

C2C12 (mouse myoblast cells), NIH3T3 (mouse fibroblast cells), SHYS5Y (neuroblastoma cells) and Hela (cervical cancer) cells were maintained in high glucose Dulbecco’s Modified Eagle Medium (DMEM) (Himedia) supplemented with 10% fetal bovine serum (FBS) (Himedia) (20% for C2C12 cells), 1X Antibacterial-Antimycotic (Himedia) solution and 1X Glutamax (Invitrogen). C2C12 and SHYS5Y were generously donated by Dr Jyotsna Dhawan, InStem and Prof Dulal Panda, IIT Bombay, respectively. HUVEC cells purchased from Lonza were maintained in EGM2-bulletkit media supplied by Lonza. Umbilical cord mesenchymal stem cells (UChMSC) were maintained in low glucose Dulbecco’s Modified Eagle Medium (DMEM) (Himedia) supplemented with 16% fetal bovine serum (FBS) (Himdeia), 1X Antibacterial-Antimycotic (Himedia) solution and 1X Glutamax (Invitrogen). UChMSC were a kind gift from Prof Jayesh Bellare, IIT Bombay. Cells were passaged at 70-80% confluency using 0.5% Trypsin-EDTA (Invitrogen) for 3 min at 37°C to detach the cells and centrifuged at 1000 rpm for 5 min. Cell were seeded on the substrates or for maintenance. The bead embedded substrates were seeded with 9000 cells/cm^2^ in 100 µl of complete media and incubated at 37°C in a humidified CO_2_ incubator for 2 h. After cell attachment, the substrates were flooded with fresh complete DMEM media and incubated at 37°C in a humidified CO_2_ incubator for overnight.

For varying seeding density studies, 500 cells/cm^2^, 3000 cells/cm^2^, 9000 cells/cm^2^ were seeded on the substrates.

### Time-lapse microscopy

The time-lapse microscopy was performed using Evos FL-Auto microscope (Life Technologies) attached with controlled environment on-stage incubator. Time-lapse imaging was started after 2 h of cell seeding and continued for 48 h duration with 30 min time interval between two consecutive images.

Cell migration was tracked using Manual tracking plugin of Image J.

### Traction Force microscopy

The PAA gel substrates with uniform stiffness were prepared by adding 100 µl of pre-polymerizing solution on 22×22 mm hydrophilic coverslip and was sandwiched by hydrophobic slide. 20 µl of PAA gel solution of same stiffness mixed with the red fluorescent beads of diameter 1 µm (Fluka) in the ratio of 50:1, along with APS and TEMED was placed on hydrophobic coverslip, then the solidified uniform gel was inverted on top of it to get the thin top layer of fluorescent beads. After 24 h of cell seeding, phase contrast and RFP fluorescent images were captured which will give the cell boundary and stressed images required for analysis, respectively. At the end, cells were killed using 200 µl 2% Triton-X without disturbing the plate and RFP fluorescent images were captured which gave the unstressed bead images. The traction force was calculated using the MATLAB code^30^ which requires constrained, unconstrained and cell boundary images as input.

### Inhibitor studies

Blebbistatin (Sigma), myosin-IIA inhibitor was used at concentration of 20 µM. Nocodazole (Sigma), microtubule polymerization inhibitor, was used at a concentration of 0.25 µM. Lysophosphatidic acid (LPA) (Sigma), phospholipid was used at concentration of 30 µM. For inhibitor studies, cells were treated with inhibitors after 2 h of seeding on substrate for 24 h.

### Image Analysis

Alignment of the nucleus is same as the alignment of the cell (Fig. S2). The images of nucleus stained with Hoechst 33342 were used to measure the alignment of the cell with respect to the center of the bead. The angle theta between the major axis of a nucleus and the radial vector starting from the epicenter of the bead and intersecting the nucleus was used as the measure of radial alignment. We defined the cells with angle θ between 0⁰ ± 45⁰ as aligned (A) (shaded region) and 90⁰ ± 45⁰ as non-aligned (N) (Fig. 1J). The alignment index (AI) is calculated using the formula AI=(A-N)/(A+N), as shown in Fig. 1J.

Analysis of cell alignment from series of time-lapse images was done using FibrilTool macro of Image J software, as reported prviously^31^. Phase-contrast images were used to determine the alignment of cells. The FibrilTool macro was customized for the analysis of time-lapse images. In brief, a square is drawn with 200 µm side and analyzed the cell alignment in that box over time. FibrilTool gives the average orientation angle of cells (θ1) with respect to x-axis of image. Another line from the center of the bead cutting the square is drawn making angle (θ2) with x-axis of the image. The acute angle between the two orientation lines is reported as the alignment of cells with respect to the embedded rigid bead. The 2kPa_B substrate showed the best alignment using the methods shown in Fig. 1I and J, therefore we calculated the value of orientation angles on 2 kPa_B at a distance <800 µm from bead periphery. The range of orientation angles of the cells was 8.7 ± 7.1. Therefore, we defined that if the orientation angle <15°, they were aligned.

Color maps of orientation angles of cells from phase-contrast images were obtained using OrientationJ Analysis plugin in ImageJ (NIH, USA) as written by Daniel Sage. The phase-contrast images in Fig. 1C-E were used as input images. OrientationJ was run with the following set of parameter for obtaining the color orientation survey for Fig. 1F and H: hue=orientation; saturation=coherency; brightness=original-image.

### Atomic Force Microscopy

The substrate was kept submerged in DPBS during measurement. The bead edge was observed under microscope and the stage was moved till it was 1500 µm away from the edge of the bead (Fig. 8B). At this point first stiffness reading was noted using TR800PB silicon nitride pyramidal tip probe on MFP-3D (Asylum Research) under contact mode force, as shown in Fig. 8A. The spring constant of cantilever ranged from 0.09 - 0.27 Nm^−1^ and frequency from 17 – 28 kHz. The force curves were then fitted with the Hertz model provided within the software from Asylum research after setting the correct parameters for tip geometry, material of the probe and the Poisson’s ratio of the material. After first reading, the stage was moved 250 µm down in y-axis direction and reading was noted. This was repeated till the bead edge was reached. From here, the stage was moved 100 µm down in y-axis direction till the other end of the bead. Then again, the gap was of 250 µm between the two readings.

**Fig. 8.**
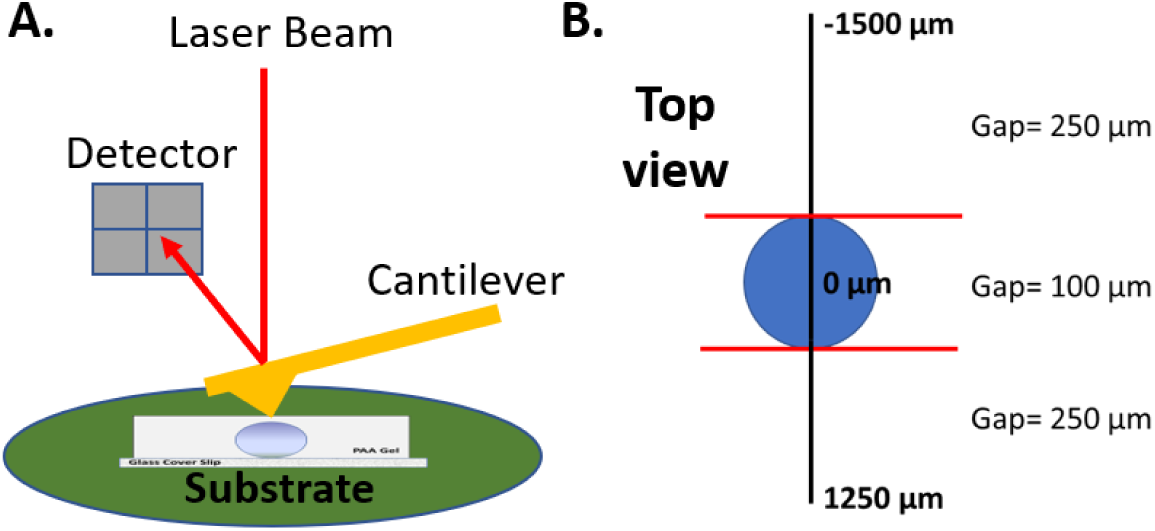
Atomic force microscopy (AFM) measurement. (**A**) Schematic of AFM probing the substrate and measuring the surface stiffness. (**B**) Top view of measurement done using AFM. Stiffness was measured at point starting from 1500 µm from the edge of the bead with 250 µm gap till the 1250 µm below the bead placed. Between the two edges of the bead the gap was of 100 µm.

### Collagen Staining

The 50 µg/ml collagen coated 2 kPa gels with and without embedded bead substrates were washed with DPBS and fixed with 4% paraformaldehyde (PFA) (pH=7.2) (Merck) for 20 min at RT followed by washing with DPBS for 3 times. The substrates were then blocked with 2% BSA in PBS (pH=7.2) (Sigma) for 2 h at RT. These gels were incubated in custom made anti-collagen antibody/serum (gifted by Prof Shamik Sen, IIT Bombay) (1:50) raised in rabbit for 4 h at RT. After incubation, gels were washed with DPBS twice, 10 min each and incubated with Alexa fluor 488 goat anti-rabbit (1:500) secondary antibody for 3 h at RT. Images were captured using LSM Carl Zeiss Confocal microscope (IITB central facility).

### Finite Element Analysis

A rectangular slab of PAA gel attached to a thin glass plate and containing an embedded glass bead at the center is modelled. The gel thickness above the top of the glass bead is denoted by t. A typical Finite Element Mesh of half the model cut by an X-section passing through the center of the glass bead is shown in Fig. 9. The in-plane dimension of the domain is 8000 x 8000 μm^2^. The thickness of the glass cover plate is taken to be 500 μm. The glass bead is spherical with a diameter of 1000 μm. Three-dimensional linear hexahedral elements C3D8 available in ABAQUS are used to discretize the domain. Several Finite Element Models are made with various values of t = 100, 150, 200 and 1000 μm. The thickness of the PAA gel ranges from 1100 μm to 2000 μm for model with t = 100 μm to that with t = 1000 μm. The total number of finite elements range from 103168 to 160768 for model with t = 100 μm to that with t = 1000 μm. As can be seen from Fig. 9, a structured mesh is created. Care was taken to ensure that the top surface of the PAA gel is meshed uniformly with equal element size of 100 μm in the three directions. The boundary conditions are such that the bottom surface of the glass plate is kept fixed. Thus, zero displacements are enforced on each node on the bottom surface in all three directions. Traction free boundary conditions are imposed on all other boundaries. Both PAA gel and glass are assumed to be isotropic and linear elastic solid materials. The Young’s modulus and Poisson’s ratio values assigned to PAA gel and glass are 2 kPa, 0.457, and 69 GPa and 0.24, respectively. In one set of simulations, perfect bonding between the glass bead and PAA gel is considered. Thus, linear elastic simulations suffice for this set. In another set of simulations, the interface between the glass bead and the PAA gel is considered to be unbonded and frictionless by specifying a clearance of 1 micron between them. Thus, non-linear finite element analyses involving contact nonlinearity at the interface in performed for this set. The bonding between the PAA gel and glass plate is perfect in all cases. The objective of the finite element simulations is to investigate the effect of embedded glass bead on the mechanics of the PAA gel substrate, which in turn might influence the patterning behavior of the cells. In particular, we are interested in calculating the apparent stiffness of the top surface of the PAA gel substrate as a function of the distance across the embedded glass bead. For this purpose, a set of finite element nodes lying on a path across the glass bead on the top surface of the PAA gel is considered as shown by red line in Fig. 9. A displacement of 20 μm is then applied on a node on this path in either Y or Z direction, and the static equilibrium problem is solved in ABAQUS with this loading to obtain the corresponding reaction force in Y or Z direction. The reaction force is divided by the imposed displacement to obtain the apparent stiffness of that point in Y or Z direction. This is referred to as apparent tangential or normal stiffness, respectively. This process is repeated for several nodes on the selected path. When plotted with distance along the path from the bead center, we get a variation of apparent tangential stiffness/normal stiffness of the top surface of the PAA gel across the bead.

**Fig. 9.**
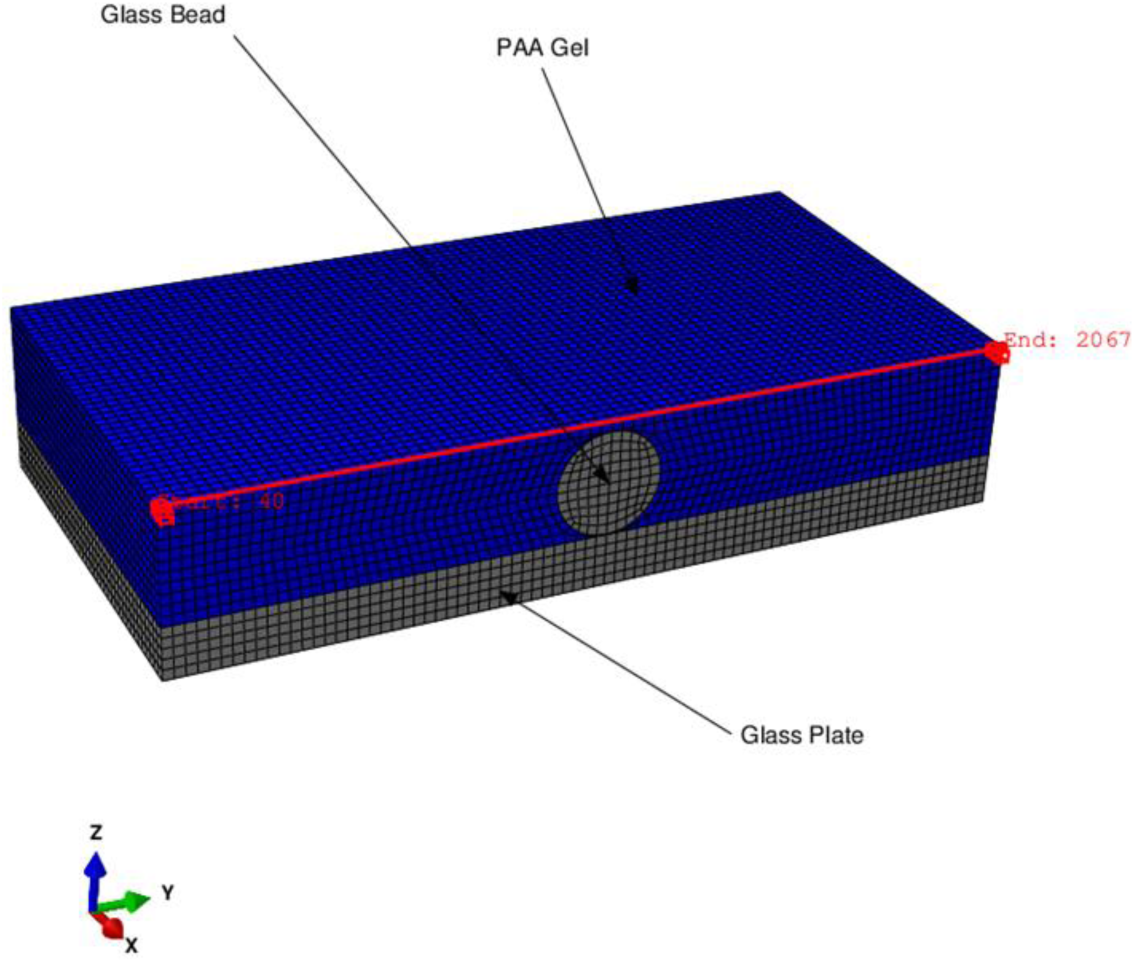
A typical Finite Element mesh of half model.

## Competing interest

The authors declare no competing interests.

## Acknowledgement

This work was supported by Wellcome Trust-DBT India Alliance (Project #IA/E/11/1/500419), IITB Seed Grant (14IRCCSG002) and Department of Chemical Engineering, IIT Bombay. We thank Bio-AFM central facility, IIT Bombay. We thank Dr. James P Butler (Harvard Medical School, Department of Medicine, Boston) for his TFM codes used for the analysis. We thank Dr. Jyotsna Dhawan, Prof Jayesh Bellare, Prof Dulal Panda, Prof Ganesh Viswanathan for generously donating the C2C12, hMSC, SHSY5Y and HeLa cells, respectively.

