## Supplementary Materials for "Asamataxis: A cooperative relayed migration in response to subsurface inhomogeneity leads to long-range self-patterning of cells"

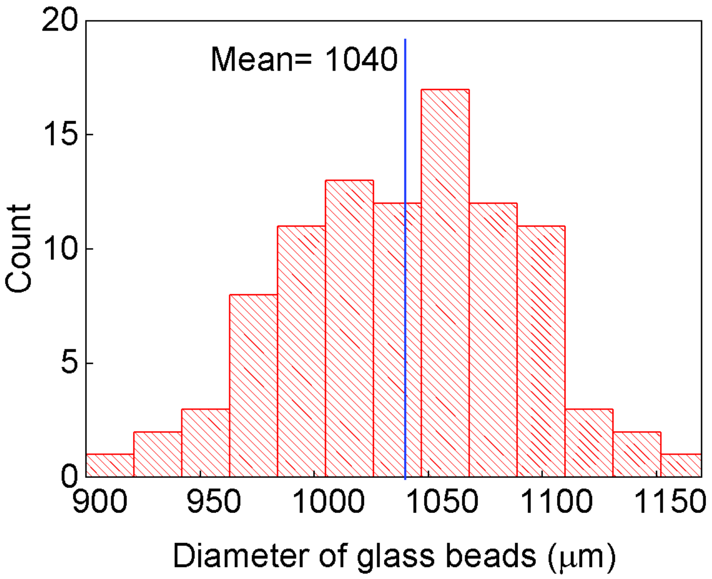


**Figure S1: Size distribution of glass beads.** The phase contrast images were captured, and the diameter of glass bead was measure using Image J. The diameter of glass bead is 1040±51 μm. (n=96 beads).

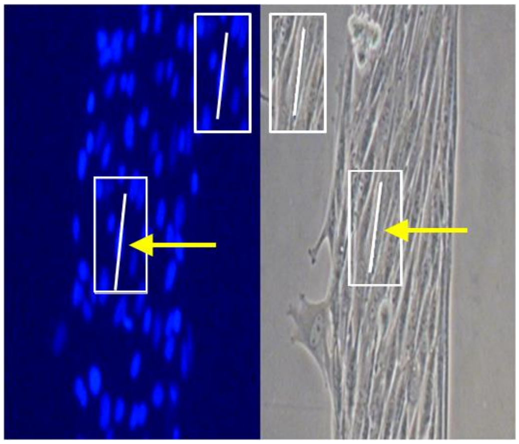

**Figure S2: Alignment between a cell and its nucleus.** The line drawn along the major axis of cell and nucleus have the same orientation with respect to the x-axis of the image. The arrowed cell and its nucleus are aligned with each other. Therefore, we can use nuclear alignment instead of cell alignment as it is challenging to identify the cell boundary.

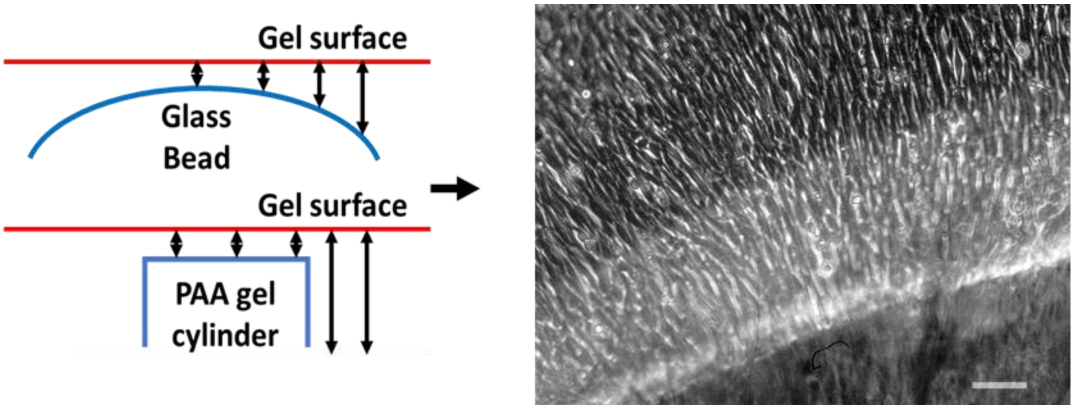


**Figure S3:** The curvature of the bead which induces thickness gradient is unimportant for radial cell alignment.

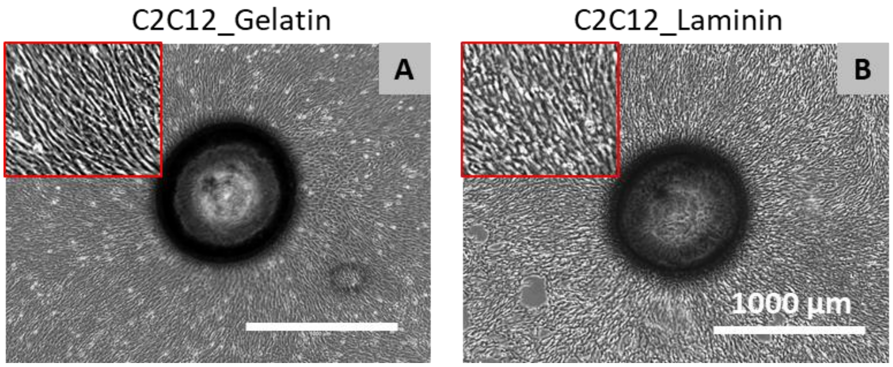


**Figure S4: Extracellular matrix independent radial alignment of C2C12 cell on 2 kPa_B substrate.** C2C12 cells were radially aligned on 2 kPa_B substrate coated with 0.5% gelatin (A) and 25 µg/ml laminin (B).

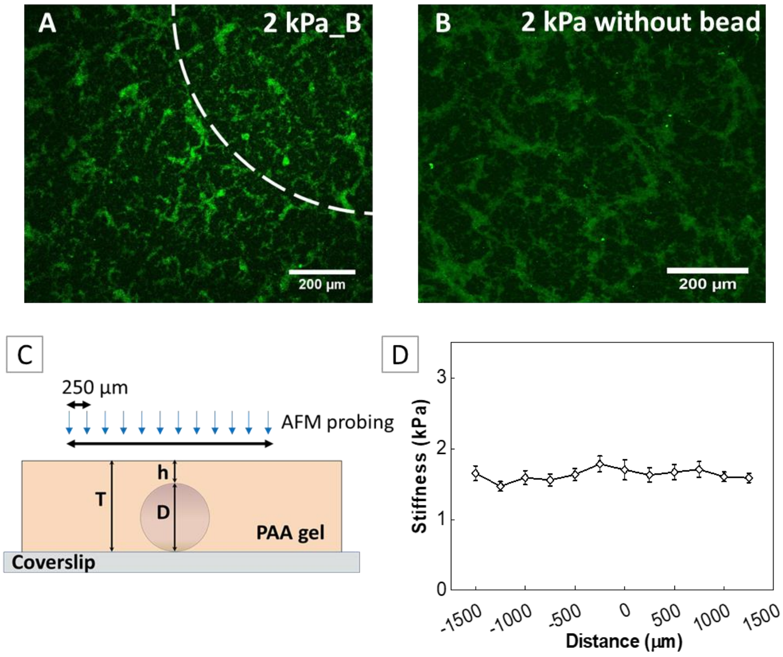


**Figure S5: Uniform coating of collagen**: Collagen staining of 2 kPa with embedded bead substrate (A) and without bead substrate (B). The collagen concentration used for coating was 50 µg/ml. The white dotted line in (A) is the periphery of the embedded bead.

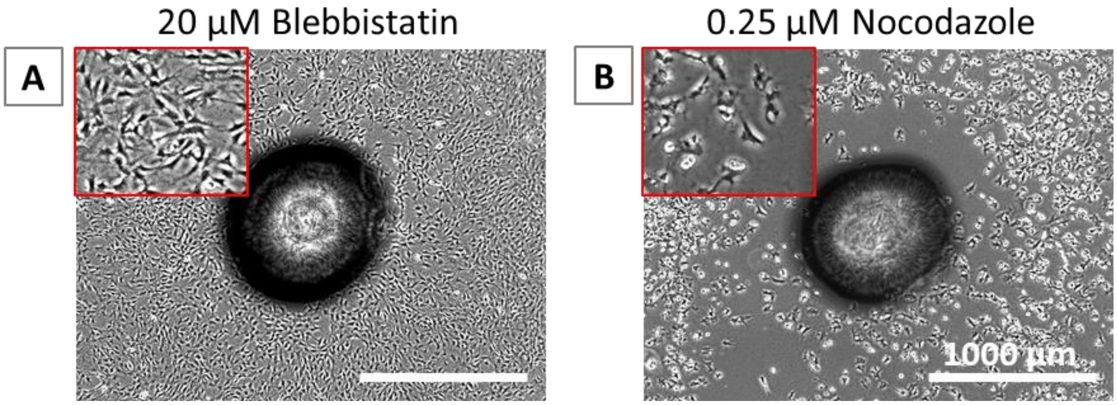


**Figure S6:** C2C12 patterning on 2 kPa_B substrate treated with (A) 20 μM blebbistatin and (B) 0.25 μM nocodazole

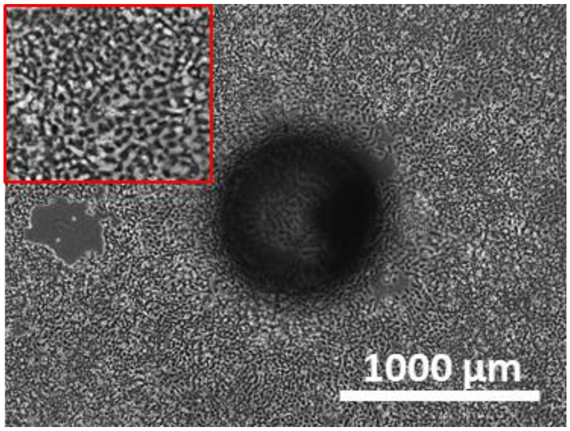


**Figure S7:** Alignment of HeLa cell on 1 kPa_B substrate coated with 50 μg/ml collagen

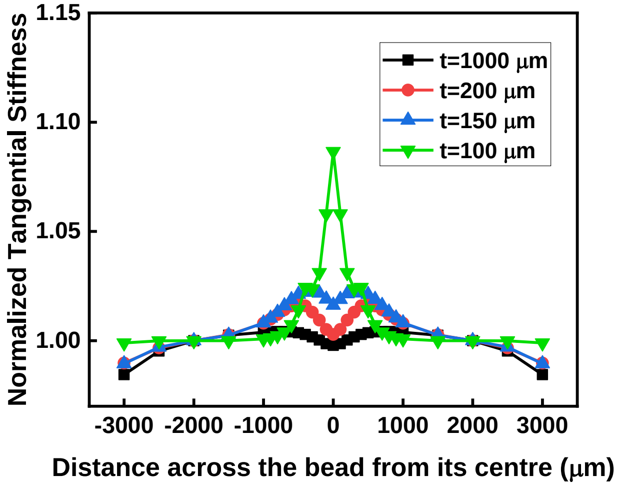


**Figure S8:** Finite Element Analysis of 2 kPa_B substrate with varying ‘t’ by applying tangential deformation at the nodes across the bead to measure the apparent increase in stiffness near embedded bead.

**Table S1:** Traction stress and morphometric parameters of different cell types used to calculate the value of $'f'$.

| Cells/conditions | Traction stress (Pa) | AR | Circularity | (1-cirularity)(AR)(TF)/stiffness |
|---|---|---|---|---|
| hMSC | 419.6 ± 82.7 | 5.7 ± 2.4 | 0.23 ± 0.11 | 919.69 |
| 3T3 | 285.5 ± 144.3 | 5.5 ± 1.6 | 0.26 ± 0.11 | 577.46 |
| Neuro | 74.2 ± 37.2 | 9.6 ± 2.9 | 0.12 ± 0.04 | 315.29 |
| C2C12 | 301.9 ± 123.7 | 2.8 ± 0.5 | 0.37 ± 0.09 | 267.25 |
| HUVEC | 292.3 ± 102.2 | 3.0 ± 1.2 | 0.53 ± 0.12 | 207.01 |
| Hela | 140.4 ± 90.9 | 2.5 ± 1.0 | 0.46 ± 0.12 | 95.22 |
| C2C12_2 kPa_Bleb | 33.2 ± 22.6 | 2.8 ± 1.2 | 0.07 ± 0.03 | 76.04 |
| C2C12_2 kPa_LPA | 343.9 ± 115.7 | 2.4 ± 0.9 | 0.32 ± 0.13 | 276.77 |
| C2C12_2 kPa_Nocod | 434.1 ± 184.5 | 1.8 ± 0.5 | 0.54 ± 0.14 | 185.32 |
| C2C12_5 kPa_control | 386.6 ± 234 | 3.2 ± 1.6 | 0.28 ± 0.12 | 178.78 |

**Video S1:** C2C12 cell migration on 2 kPa_B substrate (H = 1.2 mm, t = 0.2 mm, D = 1 mm). The initial seeding density of cells was 9000 cells/cm$^2$. Cells migrate towards embedded bead.

**Video S2:** C2C12 cell migration on 20 kPa_B substrate (H = 1.2 mm, t = 0.2 mm, D = 1 mm). The initial seeding density of cells was 9000 cells/cm$^2$. Cells are migrating randomly.

**Video S3:** C2C12 cell migration on 2 kPa_B substrate (H = 1.2 mm, t = 0.2 mm, D = 1 mm). The initial seeding of cells was 500 cells/cm$^2$. Cells are migrating randomly.

**Video S4:** C2C12 cell migration on 2 kPa_B substrate (H = 1.2 mm, t = 0.2 mm, D = 1 mm). The initial seeding of cells was 3000 cells/cm$^2$. Cells migrate towards embedded bead.
